# Evidence that APP gene copy number changes reflect recombinant vector contamination

**DOI:** 10.1101/706788

**Authors:** Junho Kim, Boxun Zhao, August Yue Huang, Michael B. Miller, Michael A. Lodato, Christopher A. Walsh, Eunjung Alice Lee

**Affiliations:** Division of Genetics and Genomics, Manton Center for Orphan Disease, Boston Children’s Hospital, Boston, MA, USA; Department of Pediatrics, Harvard Medical School, Boston, MA, USA; and Broad Institute of MIT and Harvard, Cambridge, MA, USA; Howard Hughes Medical Institute, Boston Children’s Hospital, Boston, MA, USA and Department of Neurology, Harvard Medical School, Boston, MA, USA; Department of Pathology, Brigham and Women’s Hospital, Harvard Medical School, Boston, MA, USA; Department of Cell, Molecular, and Cancer Biology, University of Massachusetts Medical School, Worcester, MA, USA

## Abstract

Mutations that occur in cells of the body, called somatic mutations, cause human diseases including cancer and some neurological disorders^1^. In a recent study published in Nature, Lee et al.^2^ (hereafter “the Lee study”) reported somatic copy number gains of the *APP* gene, a known risk locus of Alzheimer’s disease (AD), in the neurons of AD-patients and controls (69% *vs* 25% of neurons with at least one *APP* copy gain on average). The authors argue that the mechanism of these copy number gains was somatic integration of *APP* mRNA into the genome, creating what they called genomic cDNA (gencDNA). We reanalyzed the data from the Lee study, revealing evidence that *APP* gencDNA originates mainly from contamination by exogenous *APP* recombinant vectors, rather from true somatic retrotransposition of endogenous *APP*. Our reanalysis of two recent whole exome sequencing (WES) datasets—one by the authors of the Lee study^3^ and the other by Park et al.^4^—revealed that reads claimed to support *APP* gencDNA in AD samples resulted from contamination by PCR products and mRNA, respectively. Lastly, we present our own single-cell whole genome sequencing (scWGS) data that show no evidence for somatic *APP* retrotransposition in AD neurons or in neurons from normal individuals of various ages.

---

We examined the original *APP*-targeted sequencing data from the Lee study to investigate sequence features of *APP* retrotransposition. These expected features included (a) reads spanning two adjacent *APP* exons without intervening intron sequence, which would indicate processed *APP* mRNA, and (b) clipped reads, which are reads spanning the source *APP* and new genomic insertion sites, thus manifesting partial alignment to both the source and target site (Extended Data Fig. 1a). The first feature is the hallmark of retrogene or pseudogene insertions, and the second is the hallmark of RNA-mediated insertions of all kinds of retroelements, including retrogenes as well as LINE1 elements. We indeed observed multiple reads spanning two adjacent *APP* exons without the intron; however, we could not find any reads spanning the source *APP* and a target insertion site. Surprisingly, we found multiple clipped reads at both ends of the *APP* coding sequence (CDS) containing the multiple cloning site of the pGEM-T Easy Vector (Promega), which indicates external contamination of the sequencing library by a recombinant vector carrying an insert of *APP* coding sequence (Fig. 1a). The *APP* vector we found here was not used in the Lee study, but rather had been used in the same laboratory when first reporting genomic *APP* mosaicism^5^, suggesting carryover from the prior study.

**Figure 1.**
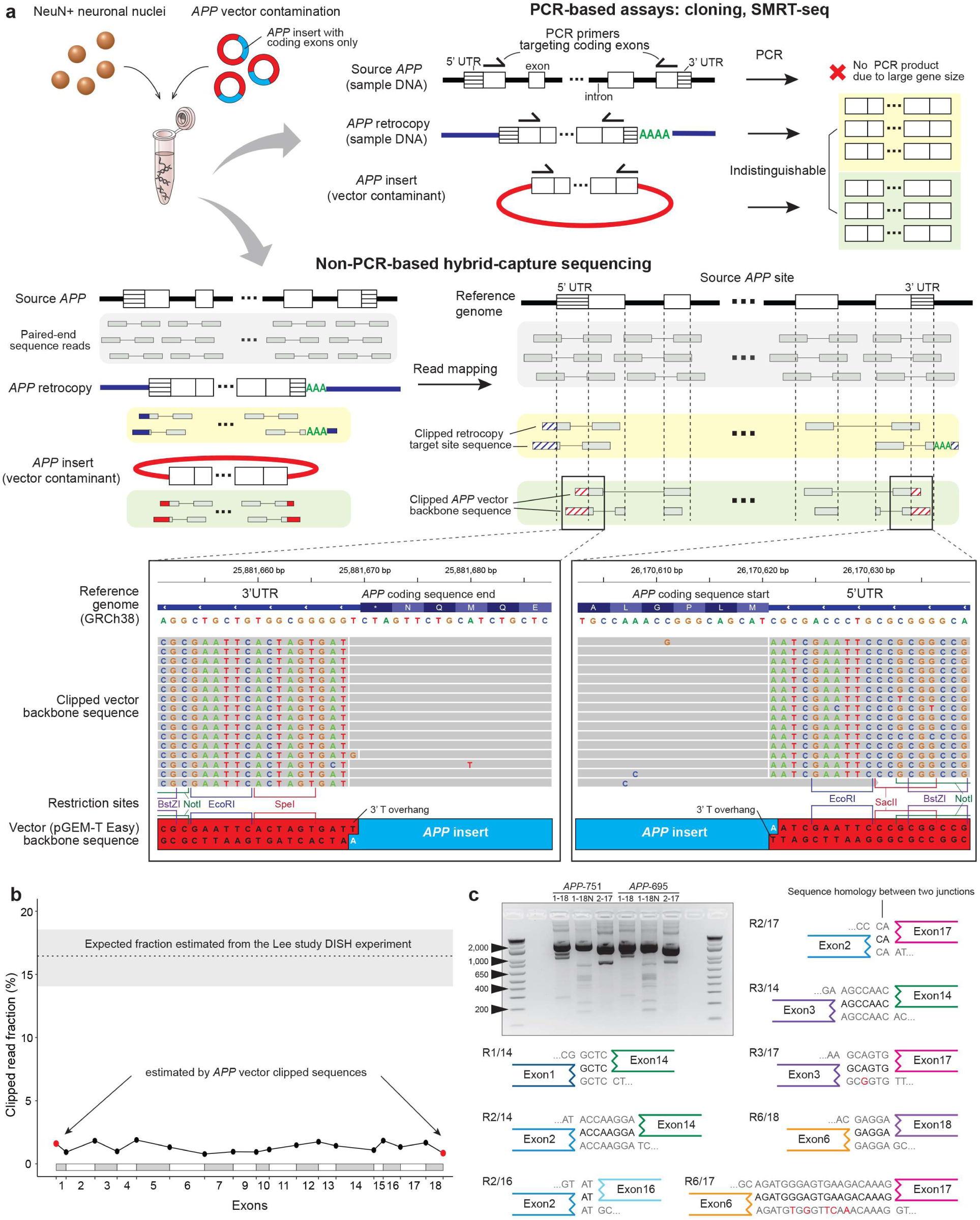
*APP* vector contamination in the Lee study. **a.** *APP* vector contamination and its manifestation in genome sequences. All designed PCR primers in the Lee study targeted only *APP* coding sequence regions shared by both *APP* retrocopy and vector *APP* insert, failing to distinguish the two sources (upper panel). In hybrid-capture sequencing, sequence reads from the flanking regions outside of the coding sequence and around the UTR regions can indicate their sources by containing the subsequence of origin (lower panel, colored in red and blue for reads originating from vectors and retrocopies, respectively). The hybrid-capture sequencing data from the Lee study clearly shows clipped reads at both ends of *APP* coding sequence with a vector backbone sequence (pGEM-T Easy), including restriction sites at the multiple cloning site, and a 3’ T-overhang (magnified panel with Integrative Genomics Viewer (IGV) screenshot). The structure of the recombinant vector contaminant and its backbone sequence are depicted, showing a perfect match to the clipped sequence. PCR duplicate reads were shown together for clear visualization of read clipping. No retrotransposition-supporting reads (blue) were detected in the hybrid-capture data. **b.** Estimated fractions of cells with *APP* gencDNA at the exon junctions in the hybrid-capture data of the Lee study. All of these exon junction fractions (black dots, fractions either from retrocopies or vector inserts) are comparable to the fraction at the coding sequence ends (red dots, fractions only from the vectors), indicating that the primary source of *APP* gencDNA is vector amplification. The dotted line on the top represents the conservative estimate of expected gencDNA-supporting ratio based on the lowest occurrence rate of *APP* retrogene insertion measured in the Lee DISH experiment (see Supplementary Methods); shaded area, 95% confidence interval. c. Electrophoresis and sequencing of PCR products from the vector *APP* inserts (*APP*-751, *APP*-695) showing novel *APP* variants as artifacts. All three PCR primer sets and three PCR enzymes used in the Lee study were tested (OneStep Ahead RT-PCR, see Extended Data Fig. 4a for other results). All novel bands were further sequenced to examine the formation of IEJs with microhomology. Eight out of 12 IEJs found both in our *APP* vector PCR sequencing and RT-PCR results from the Lee study are shown (see also Extended Data Fig. 4b). Microhomology sequences are marked with reference sequences at pre- and post-junctions (grey) and sequences derived from reads (black).

Recombinant vectors with inserts of gene coding sequences (typically without introns or untranslated regions (UTRs)) are widely used for functional gene studies. Recombinant vector contamination in next-generation sequencing is a known source of artifacts in somatic variant calling, as sequence reads from the vector insert confound those from the endogenous gene in the sample DNA^6^. We have identified multiple incidences of vector contamination in next-generation sequencing datasets from different groups, including our own laboratory (Extended Data Fig. 1b), demonstrating the risk of exposure to vector contamination. In an unrelated study on somatic copy number variation in the mouse brain^7^, from the same laboratory that authored the Lee study, we found contamination by the same human *APP* pGEM-T Easy Vector in mouse single-neuron WGS data (Extended Data Fig. 1c). We also observed another vector backbone sequence (pTripIEx2, SMART cDNA Library Construction Kit, Clontech) with an *APP* insert (Extended Data Fig. 1c, magnified panel) in the same mouse genome dataset, indicating repeated contamination by multiple types of recombinant vectors in the laboratory.

PCR-based experiments with primers targeting the *APP* coding sequence (e.g., Sanger sequencing and SMRT sequencing) are unable to distinguish *APP* retrocopies from vector inserts (Fig. 1a). Therefore, to definitively distinguish the three potential sources of *APP* sequencing reads (original source *APP*, retrogene copy, and vector insert), it is necessary to study non-PCR-based sequencing data (e.g., SureSelect hybrid-capture sequencing) and examine reads at both ends of the *APP* coding sequence. Such data can help to assess whether the clipped sequences map to a new insertion site or to vector backbone sequence. From the SureSelect hybrid-capture sequencing data in the Lee study, we directly measured the level of vector contamination by calculating the fraction of the total read depth at both ends of the *APP* coding sequence comprised by clipped reads containing vector backbone sequences (Fig. 1b, red dots). Similarly, we measured the clipped read fraction at each *APP* exon junction, which indicates the total amount of *APP* gencDNAs (either from *APP* retrocopies or vector inserts) (Fig. 1b, black dots). The average clipped read fraction at coding sequence ends that contained vector backbones (1.2%, red dots) was comparable to the average clipped read fraction at exon junctions (1.3%, black dots; P=0.64, Mann-Whitney U test), suggesting vector contamination as the primary source of the clipped reads across all the exon junctions. All the fractions at every junction are far below the conservative estimate of 16.5% gencDNA contribution based on the Lee study’s DISH experimental results (see Supplementary Information for more details on the discrepancy between sequencing and DISH results). It is incumbent on the authors to provide explanation for this significant inconsistency. Moreover, if the clipped reads were from endogenous retrocopies, the clipped and non-clipped reads would be expected to be of similar insert (DNA fragment) size distribution; however, we observed that in the Lee study, the clipped reads were of significantly smaller and far more homogeneous insert size distribution than the non-clipped reads that were from original source *APP*, thus demonstrating the foreign nature of the clipped reads (P < 2.2×10-16, Mann-Whitney U test; Extended Data Fig. 2a-c, see Supplementary Information). Finally, we found no direct evidence supporting the existence of true *APP* retrogene insertions, such as clipped and discordant reads near the *APP* UTR ends that mapped to a new insertion site, or clipped reads with polyA tails at the 3’ end of the UTR. All results from the hybrid-capture sequencing data suggest that the majority of *APP* gencDNA supporting reads actually originated from the *APP* vector contamination.

**Figure 2.**
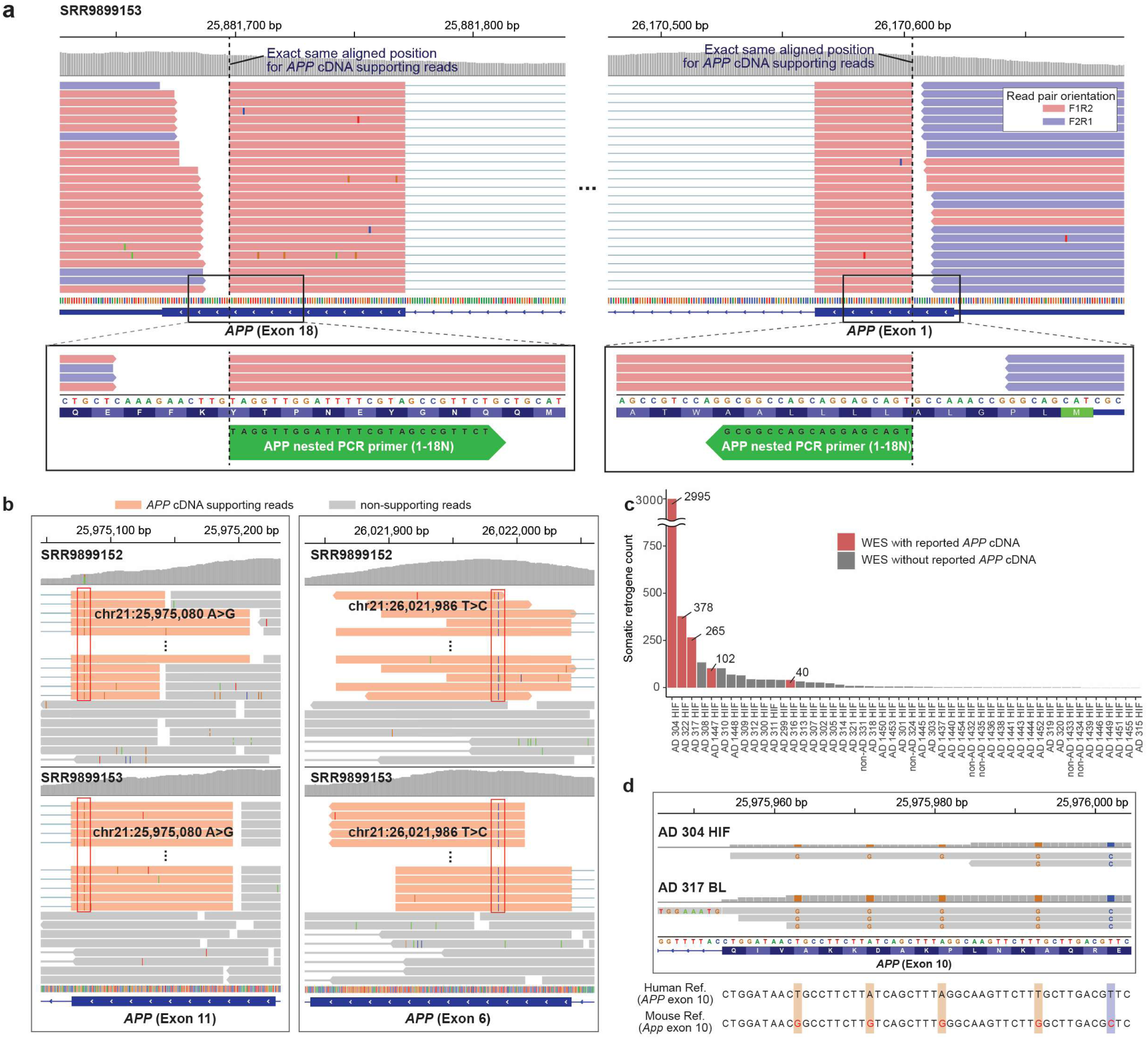
*APP* cDNA-supporting reads originate from exogenous PCR products and genome-wide RNA contamination in two recent datasets. **a.** *APP* nested PCR products found in the more recently published Lee WES dataset. Reads supporting putative *APP* cDNA are aligned to the target sites (dotted lines) of the nested PCR primers (green arrows at the bottom) used in the original Lee study. Note that a reverse complementary sequence is shown for the forward primer sequence (on the right) to show a match to the reference sequence. These cDNA-supporting reads connect exons 1 and 18 (shown with dotted lines) including an intra-exon junction (IEJ) between exons 2 and 17 (full structure not shown). **b.** Shared variants appear only in cDNA supporting reads, and appear in all of them, in the two WES datasets presented by Lee et al. (SRR989152 and SRR989153), each pooled from three AD patients. Both WES datasets (top and bottom) show the same unannotated variants at two different positions (red boxes) and only in cDNA supporting reads (orange), suggesting a common external source bearing these variants for cDNA-supporting reads. **c.** Total count of genes with potential somatic retrogene insertions in the Park et al. data. WES data with reported *APP* cDNA in the brain are marked in red. **d.** *APP* cDNA-supporting reads originating from mouse mRNA in the Park et al. data. The reference sequences of human and mouse genomes are presented together (bottom). Reads with common mismatches to the human reference sequences show mouse specific SNPs (colored bases). Clipped sequences revealed exon-exon junctions, suggesting the reads originated from mouse mRNA rather than genomic DNA. PCR duplicate reads were shown together in all IGV screenshots for clear visualization.

The authors of the Lee study have subsequently generated WES datasets from the brain samples of six AD patients and one non-AD control (SRA Accession: PRJNA558504), and reported multiple reads spanning *APP* exons without introns as evidence of somatic *APP* gencDNA^3^. We confirmed this in the data, but again, found not a single read spanning the source *APP* and any insertion sites. Instead, the data revealed anomalous patterns in a subset of reads supporting *APP* gencDNA. Those reads spanning exons 1 and 18 were aligned to the exact same start and end positions with the same read pair orientation (Fig. 2a), which is unlikely to occur in non-PCR-based exome capture sequencing. We found that the two aligned positions within exons 1 and 18 exactly match the target sites of the nested PCR primers used in the original Lee study (1-18N, Supplementary Table 1 in the Lee study). The only explanation for this observation is the contamination of the WES library by nested PCR products from the original *APP* study. This finding raises serious concerns that *APP* PCR products may also have contaminated the genomic DNA samples and were fragmented and sequenced together, generating more gencDNA-compatible reads for which we are unable to clarify the source. We also identified two unannotated single-nucleotide variants (i.e., absent in the gnomAD database^8^) in all *APP*-cDNA-supporting reads in the two independent WES libraries pooled from six AD patient samples, which is very unlikely to be observed in different individuals, thus supporting the possibility that the *APP* cDNA originated from the same external source (Fig. 2b).

An independent study by Park et al. has recently presented a small fraction of reads supporting *APP* cDNA in deep WES datasets from AD brain samples (SRA Accession: PRJNA532465; Supplementary Fig. 12 in the study)^4^. The data was free from vector contamination, but we found evidence of genome-wide mRNA (mouse mRNA in some samples) contamination predominantly in the WES datasets with reported *APP* cDNA supporting reads (Fig. 2c-d). For each AD brain sample, we counted the number of genes with potential somatic retrotransposition events by checking whether a gene had cDNA-supporting reads (i.e., reads connecting two adjacent exons skipping the intervening intron) at more than two different exon junctions in the brain sample but not in the matched blood sample from the same patient (see Supplementary Methods). All WES datasets reported by the authors to have *APP* cDNA showed an extremely high number of other genes in addition to *APP* with cDNA-supporting reads only in the brain (40-2,995 genes) (Fig. 2c). Considering that far less than one somatic retrogene insertion per sample would be expected for human cells, even for human cancers with a high rate of somatic LINE1 retrotransposition (e.g., lung and colorectal cancer)^9–11^, this result strongly suggests that cDNA-supporting reads originated from genome-wide mRNA contamination rather than from true somatic retrogene insertions. We also found some cDNA-supporting reads, including *APP* cDNA-supporting reads, originating from mouse mRNA, additionally confirming mRNA contamination of the data (Fig. 2d and Extended Data Fig. 3). Taken together, we found no evidence of genuine *APP* genomic cDNA either in the new WES data from the Lee study authors, or in the independent Park et al. data. These findings highlight pervasive exogenous contamination in next-generation sequencing experiments, even with high quality control standards, and emphasizes the need for rigorous data analysis to mitigate these significant sources of artifacts.

**Figure 3.**
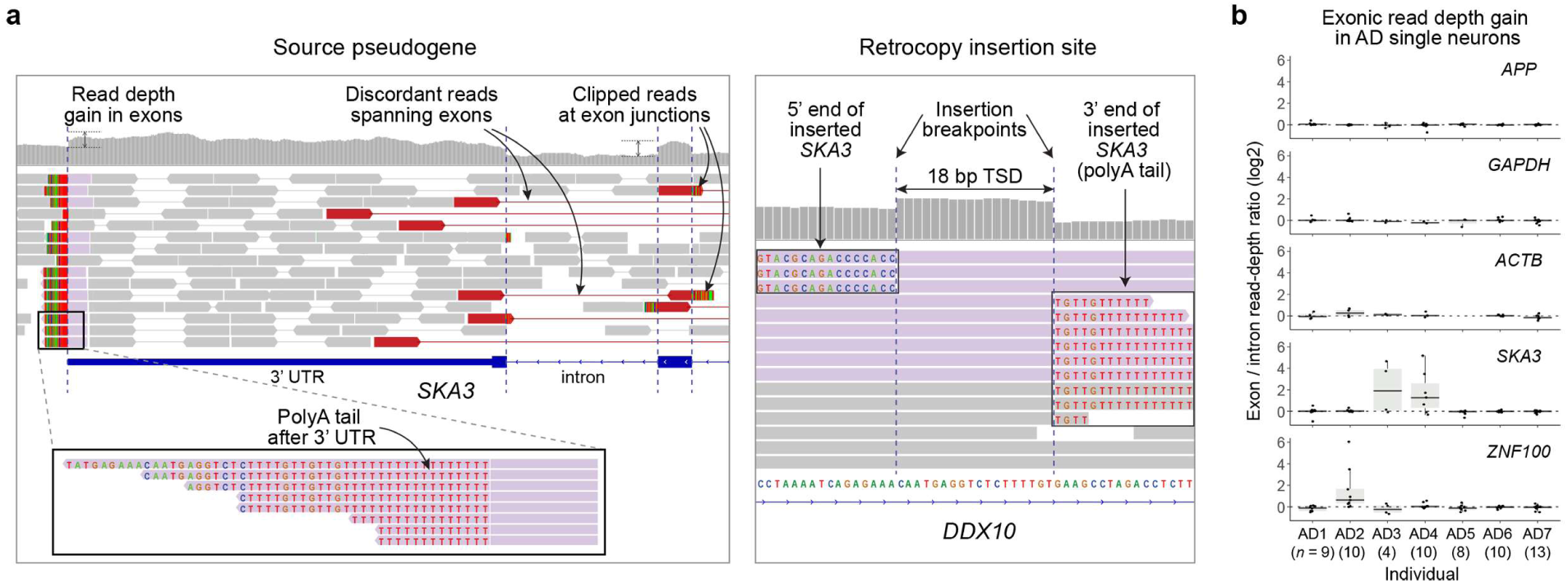
Absence of somatic *APP* retrogene insertions in our single-cell whole-genome sequencing data. **a.** A germline pseudogene insertion (*SKA3*) taken from our single-cell sequencing data. All distinctive characteristics including increased exonic read-depth, discordant reads spanning exons, clipped reads at exon junctions, 3’ poly-A tail, and target site duplication (TSD) at the insertion site are clearly observed. Mismatches including germline single-nucleotide polymorphisms and base call errors are not shown for clear visualization of insertion characteristics. **b.** No read-depth gain in *APP* exons in our AD single neurons. Each dot represents the median of exon/intron read-depth ratios across all exons of the gene in each single neuron WGS dataset from AD patients. Along with the *APP* gene, two housekeeping genes (GAPDH, ACTB) and two source genes of germline pseudogene insertions (*SKA3* in AD3 and AD4, *ZNF100* in AD2) are depicted as negative and positive controls. Single cells that had poor genomic coverage for a given gene due to locus dropout are excluded. n, number of single cells in each individual; center line, median; box limits, first and third quartiles.

The Lee study reported numerous novel forms of *APP* splice variants with intra-exon junctions (IEJs) with greater diversity in AD patients than controls. The authors also presented short sequence homology (2-20 bp) at IEJs suggesting a microhomology-mediated end-joining as a mechanism underlying IEJ formation. It is well known that microhomology can predispose to PCR artifacts^12,13^, and the Lee study performed a high number of PCR cycles in their experimental protocol (40 cycles). Thus, we tested the hypothesis that the IEJs in the Lee study could have arisen as PCR artifacts from the PCR amplification of a contaminant. To do so, we repeated in our laboratory both RT-PCR and PCR assays following the Lee study protocol using recombinant vectors with two different *APP* isoforms (*APP*-751, *APP*-695), and using the reported PCR primer sets with three different PCR enzymes as described in their study (see Supplementary Information). Indeed, with all combinations of *APP* inserts and PCR enzymes, we observed chimeric amplification bands with various sizes, clearly distinct from the original *APP* inserts (Fig. 1c, Extended Data Fig. 4a). We further sequenced these non-specific amplicons and confirmed that they contained numerous IEJs of *APP* inserts (Supplementary Table 1). 12 of 17 previously reported IEJs in the Lee study were also found from our sequencing of PCR artifacts (Fig. 1c and Extended Data Fig. 4b). Our observations suggest that the novel *APP* variants with IEJs from the Lee study might have originated from contaminants as PCR artifacts. This possibility is corroborated by the fact that IEJ-supporting reads were completely absent in the hybrid-capture sequencing data from the Lee study, and that reads supporting an IEJ in the new WES dataset by the authors originated from external nested *APP* PCR products (Fig. 2a).

To independently investigate potential *APP* gencDNA, we searched for somatic *APP* retrogene insertions in our independent scWGS data from AD patients and normal controls. Briefly, single-neuronal nuclei were isolated using NeuN staining followed by FACS sorting, whole-genome amplified using multiple displacement amplification (MDA), and finally whole-genome sequenced at 45X mean depth^14^. The dataset consists of a total of 64 scWGS datasets from 7 AD patients with Braak stage V and VI disease, along with 119 scWGS datasets from 15 unaffected control individuals, some of which have been previously published^15,16^. Our previous studies and those by other groups^14,17–19^ have successfully detected and fully validated bona fide somatic insertions of LINE1 by capturing distinct sequence features in scWGS data, demonstrating the high resolution and accuracy of scWGS-based retrotransposition detection. Therefore, if a retrogene insertion had occurred, we should have been able to observe distinct sequence features at the source retrogene site: increased exonic read-depth, read clipping at exon junctions, poly-A tail at the end of the 3’ UTR, and discordant read pairs spanning exons (Extended Data Fig. 1a). We indeed clearly captured these features at the existing germline retrogene insertions, such as the *SKA3* pseudogene insertion (Fig. 3a). If present, somatic events should be able to be detected as heterozygous germline variants in scWGS; however, our analysis revealed no evidence of somatic *APP* retrogene insertions in any of the features in any cell, not even a single *APP* gencDNA-supporting read. We also observed a clear increase in exonic read depth relative to introns for germline retrogene insertions of *SKA3* and *ZNF100* (Fig. 3b) but observed no such read depth increase for *APP* in our 64 AD and 119 normal single-neuron WGS profiles, confirming that we found no evidence of *APP* retrogene insertions in human neurons.

In summary, our analysis of the original sequencing data from the Lee study, the new WES data from the same authors, and the WES data from the independent Park study, as well as of our own scWGS data suggests that somatic *APP* retrotransposition does not frequently occur either in AD or control neurons. Rather, the reported evidence of *APP* retrocopies appears to be attributed to various types of exogenous contamination, specifically, *APP* recombinant vectors, PCR products, and genome-wide mRNA contamination. Our replication experiment also showed the possibility of PCR amplification artifacts creating spurious products that mimic *APP* gene recombination with various internal exon junctions. Thus, to support the claimed phenomenon of *APP* gencDNA, it would be necessary for the authors to present unequivocal evidence that cannot be attributed to contamination, such as reads supporting novel *APP* insertion breakpoints; however, the authors have not presented such direct evidence. In conclusion, we found no evidence of *APP* retrotransposition in the genomic data presented in the Lee study and further show that our own single-neuron WGS analysis, which directly queried the *APP* locus at single-nucleotide resolution, reveals no evidence of *APP* retrotransposition or insertion.

## Author contributions

J.K. and E.A.L. conceived and designed the study. J.K. and B.Z. designed the *APP* vector PCR and sequencing, and B.Z. performed the PCR and sequencing. M.B.M. and M.A.L. performed single-neuron sorting and sequencing. J.K. and A.Y.H. performed bioinformatic analyses. E.A.L and C.A.W supervised the study. J.K., B.Z., M.B.M., M.A.L., C.A.W., and E.A.L. wrote the manuscript.

## Competing interests

Declared none.

## Data availability

*APP* vector PCR sequences have been deposited in the NCBI Sequence Read Archive (PRJNA577966). Single-cell whole genome sequencing data of control individuals have been deposited in the NCBI Sequence Read Archive (PRJNA245456) and dbGAP (phs001485.v1.p1). Single-cell whole genome sequencing data of AD patients will be available upon request for genomic regions of *APP* and source pseudogene *SKA3* and *ZNF100*.

## Code availability

Implemented custom code for the estimation of clipped read fractions and the detection of intra-exon junctions (IEJs) is available at https://sourceforge.net/projects/somatic-app-analysis/.

## Acknowledgements

This work was supported by NIA grant K01AG051791 (E.A.L.), NINDS grant R01NS032457-20S1 (C.A.W.), NIH grant T32 HL007627 (M.B.M.), and NIH grant AG054748 (M.A.L). C.A.W. is an Investigator of the Howard Hughes Medical Institute.

---

**Extended Data Fig. 1.**
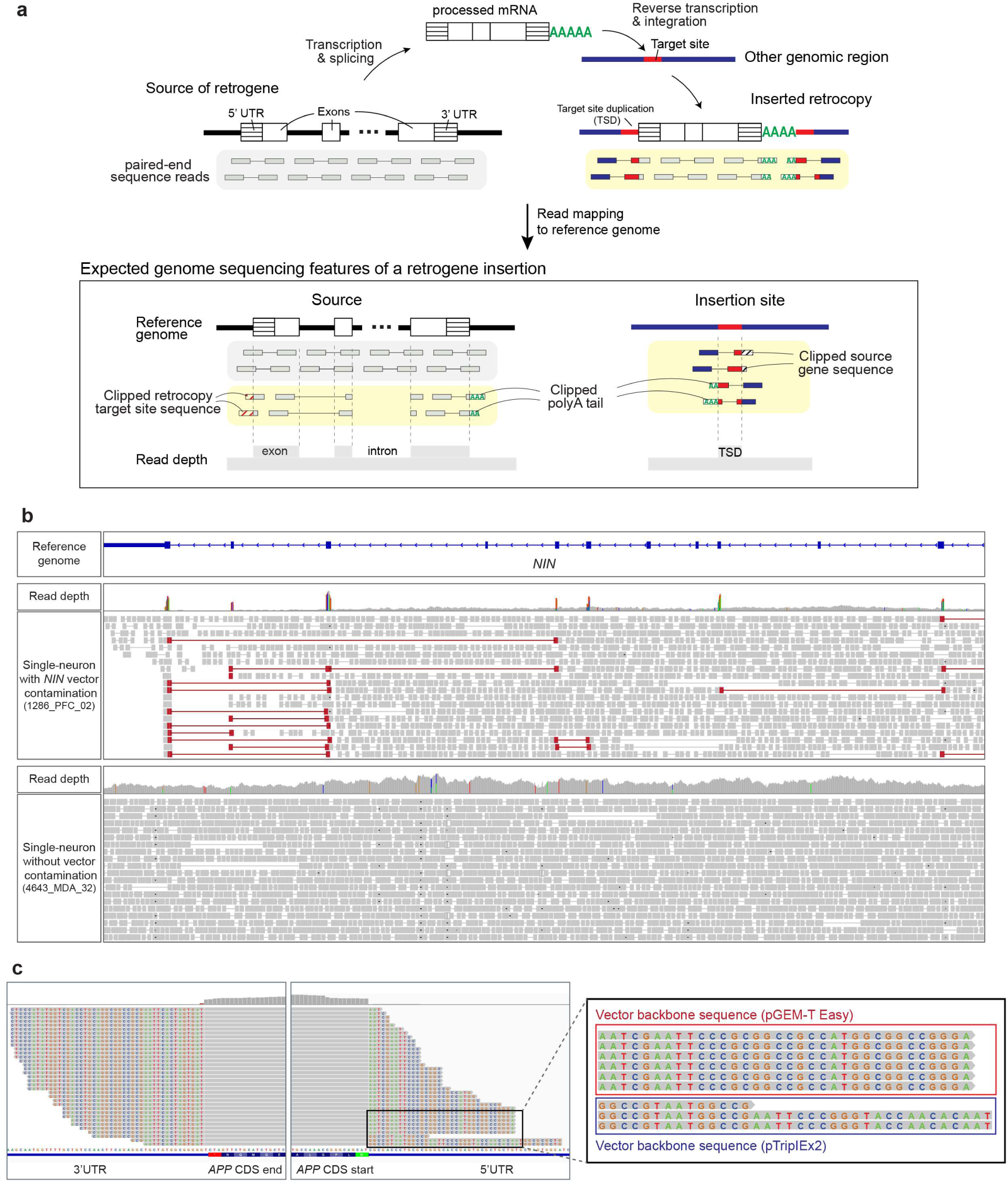
Pervasive recombinant vector contamination in next-generation sequencing. **a.** Schematic of a retrogene insertion and the characteristics expected to be captured in sequencing data: increased exonic read-depth, discordant reads spanning exons, clipped reads at exon junctions, 3’ poly-A tail, target site duplication (TSD) at the new genomic insertion site, and clipped reads spanning the retrocopy and insertion sites. Vector contaminants can mimic most characteristics of true retrogene insertions, except for features related to new insertion sites and the insertional mechanism such as polyA tail and TSD, since recombinant vectors contain inserts of processed gene-coding sequences. **b.** Recombinant vector contamination from an experiment performed in the Walsh laboratory. Four single human neurons (1286_PFC_02, 1762_PFC_04, 5379_PFC_01, 5416_PFC_06) in our previous publication contained contamination by sequences from a mouse Nin recombinant vector^20^. The homologous human gene region of the source gene (NIN) is visualized by the IGV browser for a vector contaminated cell (upper panel) and an unaffected control cell (lower panel). Contamination characteristics including increased exonic read-depth and discordant reads spanning exons (reads colored in red) were clearly identified. Note that because the contaminant inserts were derived from the mouse Nin gene and mapped here on the human reference genome, numerous mismatches were observed in exonic regions (indicated by colored vertical bars in the read depth track). **c.** Another *APP* vector-contaminated dataset from the Chun laboratory^7^. This mouse single-neuron WGS data was contaminated by the same *APP* recombinant vector detected in the Lee study^2^. An additional *APP* plasmid vector was also identified in this experiment (magnified panel), suggesting contamination by multiple recombinant *APP* vectors in the laboratory.

**Extended Data Fig. 2.**
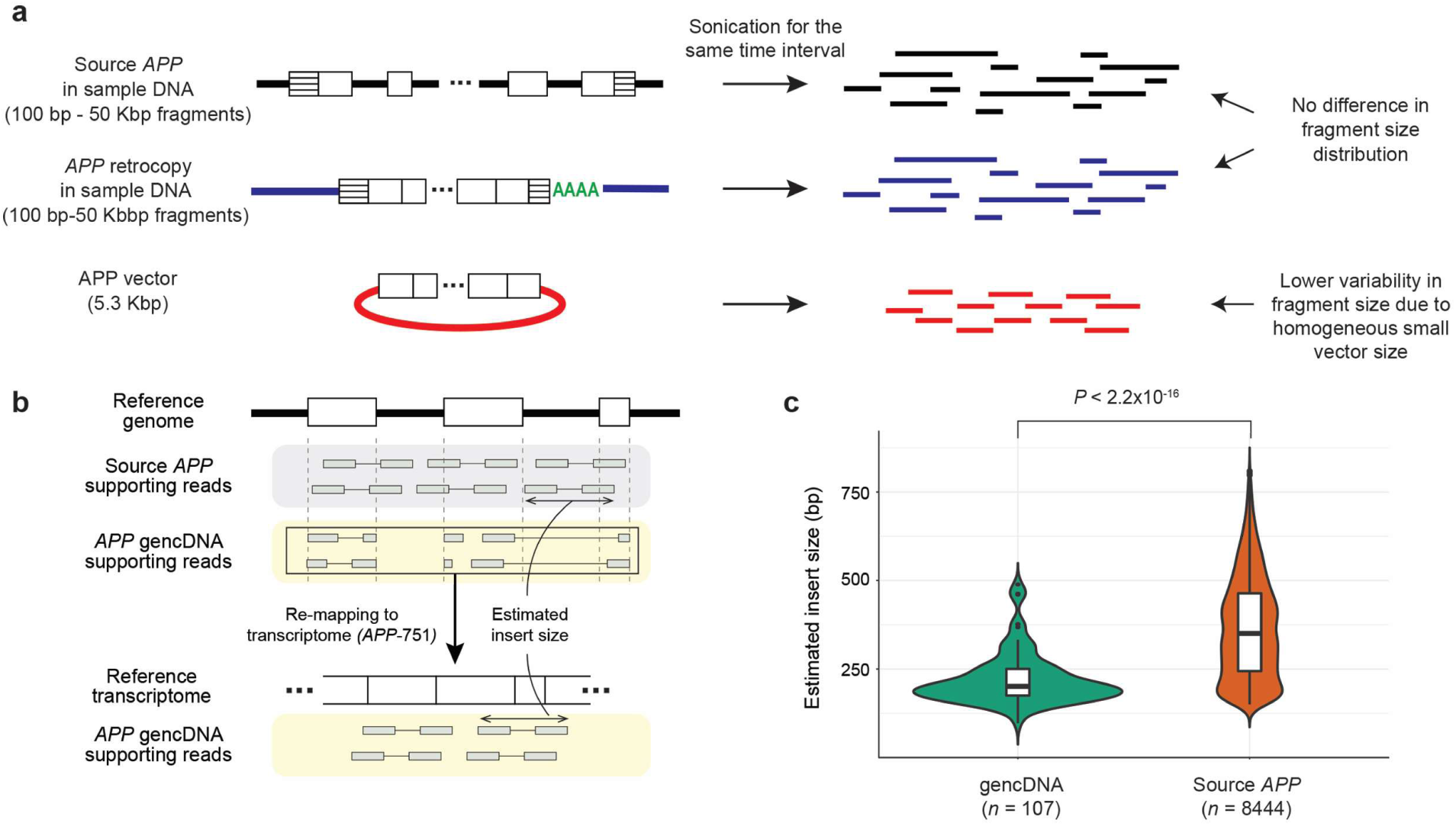
Evidence that recombinant vector contamination is the major source of *APP* gencDNA. **a.** Schematic of the DNA fragment size distribution for each *APP* source (source *APP*, *APP* retrocopy, *APP* vector). Fragments from *APP* vectors are expected to be more homogeneous and smaller in size than those from other sources due to the fixed and relatively small vector size. **b.** DNA fragment (or insert) size estimation. Sequence reads mapped to *APP* exon junctions were divided into two groups: source *APP* (reads containing intron sequences) and *APP* gencDNA (reads clipped at the exon junction) supporting reads. gencDNA supporting reads were remapped to the *APP* reference transcript sequence (*APP*-751) to estimate insert sizes. **c.** Comparison of insert size distribution between source and gencDNA supporting reads. n, number of read pairs in each group.

**Extended Data Fig. 3.**
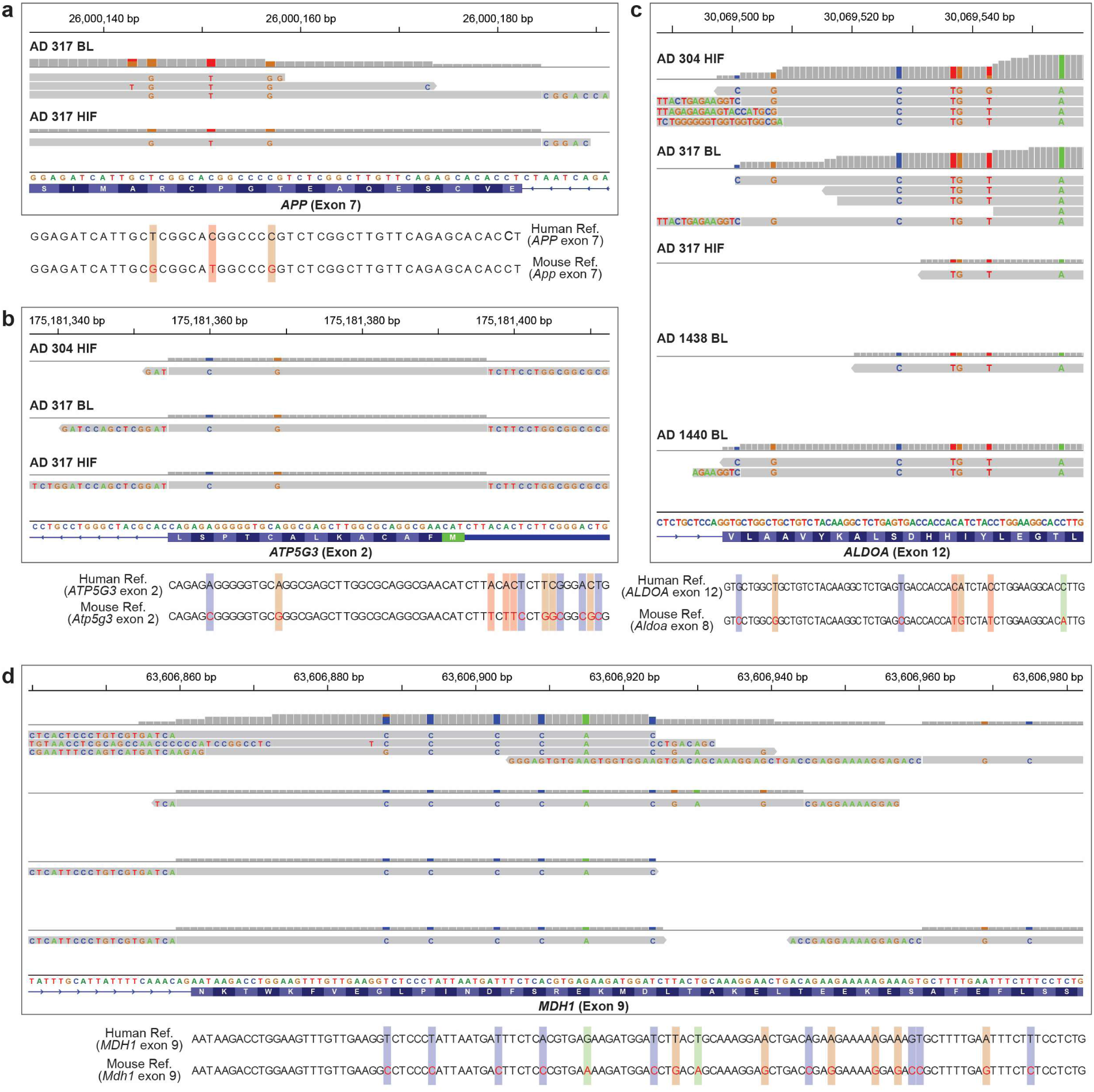
Mouse mRNA contamination in the Park et al. data. cDNA-supporting reads with mouse-specific SNPs identified in multiple samples are presented. Clipped sequences at the exon junction are not matched to the intron but rather are matched to the adjacent exon, indicating the reads originated from mouse mRNA rather than from genomic DNA. Some read clipping occurs slightly off the exon junction (typically 2-3 bp) due to the sequence homology of splicing donor/acceptor sites.

**Extended Data Fig. 4.**
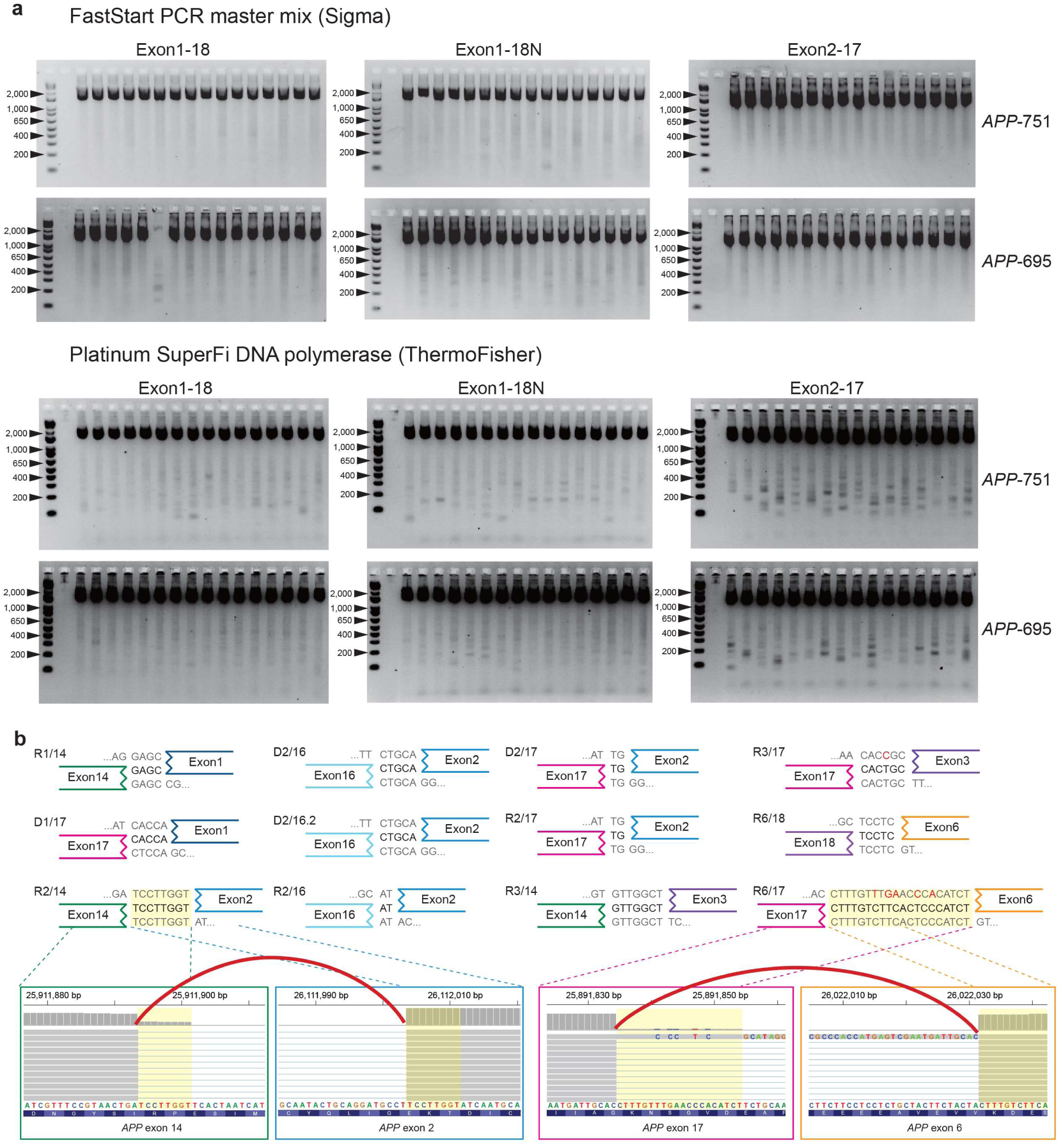
Novel *APP* variants with intra-exon junctions as PCR artifacts. **a.** Electrophoresis of PCR products from the vector *APP* inserts (*APP*-751, *APP*-695) showing novel *APP* variants as artifacts. Results of two PCR enzymes (FastStart PCR master mix, Platinum SuperFi DNA polymerase) with three primer sets are presented. All combinations generated novel bands smaller than the expected PCR product. **b.** PCR-induced IEJs with homologous sequences at each junction identified by Illumina sequencing. Twelve IEJs from our vector PCR sequencing showed exactly the same sequence homologies and genomic coordinates as IEJs reported in the Lee study. For two IEJs, IGV browser images show pre-(left) and post-junction sites (right) connected by split reads spanning the IEJ (red arc). Because IGV displays forward strand sequences of the human reference genome, all IEJ sequences were also reverse complemented for consistent visualization.

## Notes

#### Summary of Updates

Analysis results updated with additional figures (Fig.2, Extended Fig.3) to incorporate new data

