## Supplementary Information for "Evidence that APP gene copy number changes reflect recombinant vector contamination"

### Supplementary Methods

#### Detection of vector contamination in the *APP*-targeted hybrid-capture sequencing data from the Lee study[1]

We downloaded the raw Illumina paired-end reads of the Lee study *APP*-targeted hybrid-capture sequencing data from the Sequence Read Archive (SRA accession number: PRJNA493258). We trimmed TruSeq adapter sequences from the FASTQ files using Cutadapt (version 1.14)[2] and mapped the trimmed reads to the human reference genome (GRCh38). Whereas the Lee study used STAR[3] for read alignment, we used BWA-mem (version 0.7.17)[4] since it is sensitive to align partially mapped reads to the genome with more efficient read clipping than STAR. Duplicated reads from the mapped BAM file were marked using Picard (version 2.8.0), and indel realignment and base-quality recalibration were then performed using the Genome Analysis Toolkit (GATK, version 3.5)[5]. The analysis-ready BAM file was then examined for *APP* vector contamination using Vecuum (version 1.0.1)[6] and was also visually inspected using the Integrative Genomic Viewer (IGV)[7]. Both results demonstrated clear read clippings at both ends of the *APP* coding sequence rather than 5'/3' UTRs, indicating a high possibility of external contamination by *APP* recombinant vectors since recombinant vectors typically do not contain UTR sequences. We collected clipped sequences to generate a consensus sequence of the vector backbone. The constructed consensus sequence was queried against the NCBI nucleotide collection database using NCBI BLASTN[8], to identify the type of vector used. Identified vectors were further confirmed by matching the consensus sequence to vector backbone sequences from the Addgene repository[9], when available.

**Estimation of the fraction of cells with gencDNA in the hybrid-capture data from the Lee study**

*APP* gencDNA could originate from two different sources: true *APP* retrocopies or vector *APP* inserts. We first directly measured the contribution of vector contamination by calculating the vector fraction, *i.e.,* the number of clipped reads containing vector backbone sequences over the total read depth at both ends of the *APP* coding sequence (CDS; red dots in Figure 1b). We then measured the gencDNA fraction, *i.e.,* the number of clipped reads from both *APP* retrocopy and vector insert over the total read depth at each *APP* exon junction (black dots in Figure 1b). By comparing the gencDNA fractions to the vector fractions, we can estimate the proportion of gencDNA originating from true *APP* retrocopies.

In the fraction calculation, we need to consider the difference in hybridization-based capture efficiency between CDS ends and exon junctions (Supplementary Fig. 1a). Specifically, *APP* targeting probes are less efficient at capturing DNA fragments that contain long vector backbone sequences but short *APP* sequences than they are at capturing *APP* gencDNA fragments at both sides of an exon junction (Supplementary Fig. 1a). To adjust for this capture efficiency bias, in our clipped fraction calculation, we only used a paired-end read when the mate read mapped toward the inside of the exon, *i.e.,* those that could be reliably captured by the *APP*-targeting probes (Supplementary Fig. 1b, bolded reads). These corrected fractions were used to compare the amount of vector contaminant to the amount of gencDNA (Fig. 1b red dots and black dots, respectively). For each exon junction, we reported the average of the two fractions from the two adjacent exon-intron junctions (Fig. 1b, black dots). Overall, there was no difference between the vector fractions and the gencDNA fractions (1.2% vs. 1.3% on average; $P$=0.64, Mann-Whitney U test), indicating that the primary source of gencDNA was *APP* vector DNA.

**Calculation of the expected gencDNA fraction based on the Lee study DNA in situ hybridization (DISH) experiment**

The Lee study measured the copy numbers of gencDNA for each individual cell using two different DISH probes: one targeting the exon16-17 junction ($DISH_{16/17}$) and the other targeting the intra-exonic junction (IEJ) between exon 3 and 16 ($DISH_{3/16}$). They reported DISH measurements for more than a hundred cells for each of six AD patients and six normal controls (Fig.5, Extended Data Fig. 5 in the Lee study). We estimated the expected gencDNA fraction based on the Lee study DISH probe measurements (Figure 1b, dotted line). It would be ideal to be able to match the DISH results to their *APP*-targeted capture sequencing data from the same AD patients; however, the authors did not report which patients were profiled with *APP*-targeted sequencing. Therefore, we selected the AD patient (SAD4) who showed the least amount of *APP* gencDNA in their DISH experiments to create the most conservative estimate of the gencDNA fraction that would be expected to be present in the Lee study capture sequencing data to support their claim. Specifically, using the $DISH_{16/17}$ probe measurements for 155 single cells of the SAD4 patient, we calculated the expected fraction of gencDNA clipped reads in the capture sequencing, as follows:

$$F = \frac{CN_{avg}}{2 + CN_{avg}} \times 0.5$$

$CN_{avg}$ represents the average copy number measured by the DISH experiment. The two in the denominator was added since every cell contains two copies of the source *APP* gene. The fraction was then multiplied by 0.5 to account for the random chance that a clipped read spanning the exon junction would be mapped to only one side of the exon end. Compared to this conservative estimate, the gencDNA fractions that we actually observed from the capture sequencing data was far lower. This represents a notable inconsistency between the DISH and the capture sequencing experimental results in the Lee study (Fig. 1b).

**Unexplained discrepancy of APP retrocopy estimates from sequencing *vs* DISH experiments in the Lee study**

The Lee study estimated *APP* retrogene copies solely based on their DISH experiments, and the estimates were greatly inconsistent with our estimates from their sequencing data, even though the sequencing data were generated from the same individuals as the DISH experiments. Specifically, their DISH experiments for the exon 16-17 junction ($DISH_{16/17}$) showed that each AD neuron carried 1.70 copies of somatic *APP* gencDNA on average, and the AD brain SAD1 had 95.6% of neurons with 4.33 copies of gencDNA per neuron. Based on these estimates, the expected ratio of *APP*-gencDNA-supporting reads over the total read depth at the exon 16-17 junction is from 45.9% (with 1.70 copies) to 68.4% (with 4.33 copies), which are stronger signals than a germline one-copy insertion (33%). By contrast, their capture sequencing data from bulk tissue of the same individuals showed only a small fraction (2.68%) of reads supporting the exon 16-17 junction and even those we found to in fact reflect vector contamination. The DISH results for the IEJ between exon 3 and 16 ($DISH_{3/16}$) also showed a similarly high rate—an average of 1.23 copies— of *APP* gencDNA per AD neuron thus expected to have 38.1% of supporting reads over the total depth at the breakpoints, but no read supporting the IEJ was detected in their capture sequencing data. The authors have not explained this large discrepancy in their results by DISH and sequencing experiments.

**Comparison of DNA fragment, or insert size, for reads originating from the source *APP* and gencDNA**

According to the genomic DNA extraction protocol (DNeasy and QIAamp DNA Mini kits) used in the Lee study, purified DNA fragments are expected to be 100bp to 50 Kbp in size, predominantly 30 Kbp. In contrast, the *APP* vector we detected in the Lee study (*APP* gene in the pGEM-T Easy Vector) has a smaller, fixed size (5.3 Kbp). Therefore, with sonication for the same time interval in the library preparation process of hybrid-capture sequencing, vector-derived DNA fragments are likely to be smaller and be more homogeneous in size than DNA fragments from original *APP* or *APP* retrocopies (Extended Data Fig. 2a). In order to estimate DNA fragment sizes from the source *APP*, we extracted read pairs spanning *APP* exon-intron junctions (*i.e.,* those containing intron sequences) from the BAM file and measured their insert sizes based on their mapped coordinates.  Read pairs for which the insert size differed from the mean by more than three standard deviations were considered as discordant reads and discarded for conservative estimation of the insert size of original *APP* supporting reads. To estimate DNA fragments from gencDNA, we extracted read pairs clipped at the exon junctions and remapped them to the *APP* reference transcript sequence (*APP*-751; NCBI CCDS ID: CCDS33523.1) to obtain insert sizes based on the mapped coordinates (Extended Data Fig. 2b).

**Detection of *APP* vector contamination in mouse single-neuron whole-genome sequencing data from another study from the Chun laboratory**[10]

We confirmed that the vector contaminant we found in the Lee study (*APP* gene in the pGEM-T Easy vector) was not used in the corresponding work, but rather had been used in another study on genomic *APP* mosaicism[11] from the same laboratory. To verify whether this contamination has only affected the Lee study or whether there may be broader contamination, we investigated another sequencing dataset published from the same group and checked for *APP* vector contamination. We downloaded the raw single-end FASTQ files from the study of copy-number variations in single-cell whole-genome sequencing of mouse neurons[10] (SRA accession number: PRJNA415480), which is work completely unrelated to the *APP* studies. We first merged all 522 FASTQ files into one integrated FASTQ file to increase the sensitivity to vector contamination detection. Next, we trimmed Nextera adapter sequences from the integrated FASTQ file using Cutadapt. Due to sequence homology between mouse and human *APP* sequences, we mapped the reads to the human-mouse hybrid reference genome (GRCh38/mm10) rather than to the human reference genome for unambiguous read mapping. BWA-mem, PICARD, and GATK were used for read mapping, marking duplicates, indel-realignment, and base-quality recalibration as described above. Visual inspection of mapped data for the human *APP* region identified two different types of *APP* recombinant vectors, including exactly the same one as in the Lee study, suggesting contamination of multiple types of recombinant vectors in the laboratory.

**Detecting somatic retrogene candidates from the Park et al. data**

An independent study by Park et al. has reported evidence of somatic *APP* retrotransposition in AD patients from deep whole-exome sequencing data, showing *APP*-cDNA-supporting reads only in brain tissue samples but not in matched control samples [12]. We tried to identify from their data all somatic retrogene candidates, including *APP*, to check whether the data showing *APP* cDNA-supporting reads identified by Park also showed cDNA-supporting reads from various other genes, which would flag possible exogenous contamination given the rare incidence of somatic retrogene insertion in human cells[13,14].

We assumed there to be four different sources of cDNA-supporting reads: 1) somatic retrotransposition, 2) germline pseudogene insertion, 3) exogenous contamination, and 4) misalignment artifacts. We were able to eliminate the possibility of germline pseudogene insertion because we are trying to find candidates that are only found in brain tissue samples and not in matched samples. We set up criteria to identify somatic retrogene candidates as genes that have more than two distinct exon junctions supported by at least one cDNA-supporting read in the brain sample, but no cDNA at all at any exon junction in the matched control sample.

We downloaded the raw paired-end FASTQ files from Park et al. (SRA accession number: PRJNA532465), which contained whole-exome sequenced reads of 48 brain-blood matched samples and 15 unmatched brain samples. Since we tried to identify somatic candidates absent in the matched control tissue, we only analyzed the 48 matched pairs. To detect cDNA-supporting reads, we first mapped all the data to the human reference genome (GRCh38) using STAR (version 2.5.4a)[3] with the options described in the Lee study and also used in the Park study (--outSAMattributes All --outFilterScoreMinOverLread 0.8 --outSJfilterCountTotalMin 1 1 1 1). We then filtered the aligned BAM files using SAMtools (version 1.9)[15] to select only cDNA-supporting reads by excluding reads with the tag 'jI:B:i,-1', which indicates that no junction was detected.

We found that extracted cDNA-supporting reads from STAR contained numerous false positives caused by misalignment of reads or by errors in splitting reads due to sequence homology of splicing donor/acceptor sites. Therefore, we considered only uniquely mapped cDNA-supporting reads (mapping quality of 255) so as to reduce misalignment artifacts. We also checked sequence homology between split reads and the intronic sequence to which the full read would be aligned if the read split had not occurred. Reads that show high similarity (>90% identical) between the split read and the intronic sequence were considered to be erroneously split and were thus discarded. We could still find misaligned reads with higher mapping scores when we apply other alignment tools such as BLAT. To remove them, we additionally aligned all the data with BWA-mem with preprocessing steps to remove duplicates, realign indels, and recalibrate base quality; we then compared the mapping position of cDNA-supporting reads between the STAR- and the BWA-aligned BAMs. All reads with discordant mapping positions were filtered out as misalignment artifacts. Lastly, we analyzed the mapping positions of the mate read of each cDNA-supporting read and filtered out the read if the mate was aligned to the intronic sequence and therefore did not support the processed form of the gene. Although these filtering steps are quite conservative and might filter out some true cDNA-supporting reads, we still were able to find somatic retrogene candidates with cDNA at more than two different exon junctions for up to 2,995 source genes from a single sample. This serves as a clear indication of exogenous contamination of the data rather than indicating true somatic retrotransposition.

**Replication of PCR artifacts that mimic various *APP* recombinant variants**

We tested our hypothesis that *APP* variants with IEJs reported in the Lee study could have arisen as PCR mis-pairing artifacts between partially homologous sequences during the PCR amplification of *APP* vector contaminants. First, we repeated the PCR assays following the Lee study protocols using 250 pg of recombinant vectors with two different isoforms of *APP* inserts (pCAX *APP* 751 [Addgene plasmid # 30138 ; http://n2t.net/addgene:30138 ; RRID:Addgene_30138] and pCAX *APP* 695 [Addgene plasmid # 30137 ; http://n2t.net/addgene:30137 ; RRID:Addgene_30137]) as templates, respectively. All combinations of three PCR enzymes (OneStep Ahead RT-PCR (Qiagen), FastStart PCR master mix (Sigma), Platinum SuperFi DNA polymerase (ThermoFisher)) and three reported PCR primer sets (*APP* 1-18, *APP* 1-18N, *APP* 2-17; Supplementary Table 1 in the Lee study) were tested with two different final primer concentration settings (0.5 μM and 1.0 μM).

For OneStep Ahead RT-PCR, the cycling program of low annealing stringency PCR was 45 °C for 15 min; 95 °C for 5 min; 40 cycles of 95 °C for 15 sec, 55 °C for 15 sec, 68 °C for 2.5 min; 68 °C for 5 min. High annealing stringency PCR was performed for 40 cycles using the FastStart PCR master mix with the following cycle settings: 95 °C for 30 sec, 65 °C for 30 sec, and 72 °C for 2.5 min. We used the Platinum SuperFi DNA polymerase with the following cycle settings: 98 °C for 10 sec, 65 °C for 10 sec, and 72 °C for 1.5 min. All PCR products were run on 2% agarose gels. For all PCR combinations, we observed multiple chimeric amplification bands that were clearly distinct from the correct band of *APP* inserts.

**Illumina sequencing of *APP* vector PCR products and identification of IEJs with microhomology**

We further sequenced the non-specific *APP* vector PCR amplicons and confirmed the existence of IEJs. All of the chimeric amplification bands were cut and recovered from 2% agarose gels using a QIAquick Gel Extraction Kit (28706; QIAGEN). Extracted DNA was sonicated into fragments with an average size of 200 bp using Covaris S2 with the following settings: sample volume, 50 µL; water level, 12; temperature, 7°C; intensity: 5; duty cycle, 10%; cycles per burst, 200; and treatment time, 120 s. Amplicon fragments were endrepaired, dA-tailed, and adaptor-ligated using the KAPA Hyper Prep Kit (KK8503; KAPA Biosystems). Amplicon libraries for each combination were labeled with unique dual index and paired-end sequenced (2×151 bp) on the Illumina HiSeq platform at Microgen, Inc. Reads were aligned against the sequence of the *APP* reference transcript (*APP*-751) using BWA-mem.

We identified IEJs with microhomology based on read clipping information provided by BWA-mem. When a read contains an IEJ, BWA-mem first clips the read at the junction site and maps the longer remaining part to one exon. It then remaps the clipped-out subsequence to another exon and tags it as a secondary alignment with the same read name. For these primary-secondary alignment pairs, we searched the microhomology of the reference sequence shared between those two mapped regions. Specifically, we extracted 50 bp of the reference sequence from the mapped pre-junction part of the primary alignment and also from the flanking region of the secondary alignment, and compared their subsequences ranging from two to 50 bp. A subsequence pair was considered to have microhomology if the two had more than 75% concordance, the minimum concordance of microhomology that the Lee study reported (Fig. 1e in the Lee study). If a given pair contained multiple microhomology subsequences, we selected the longest one as representative and reported only that one. A total of 17,011 IEJs with microhomology were identified from six different PCR experiments. 12 of 17 previously reported IEJs in the Lee study were also detected from our sequencing of PCR artifacts, suggesting that the reported IEJs may have arisen from PCR errors. IEJ-supporting reads were extracted and realigned by STAR[3] for visualization.

**Analysis of retrogene insertions in our independent scWGS data**

We examined two different features of somatic *APP* retrogene insertions in our own scWGS data from AD patients and normal controls: 1) increased read depth in exons compared to introns of the *APP* gene and 2) reads spanning adjacent exons of the *APP* gene without an intervening intron sequence.

For each scWGS data, we measured the ratio of the exonic read depth to the intronic read depth from the *APP* gene to represent the exonic read depth gain of a single cell. Due to the uneven genome amplification in single cell sequencing, we measured the read depth not from the entire gene region, but from the small flanking regions (50 bp) of each exon-intron junction. Since short homologous sequences at junctions (*e.g.,* splice sites) often cause imprecise read clipping by BWA-mem, we discarded the read depths of the 10 bases nearest to the junction in calculating the average read depth for each exon and intron. The exon/intron read depth ratio was calculated for every exon end using these average values. If either exonic or intronic average read depth was less than one for a given exon end, that end was excluded for the calculation of the ratio. The median of all of the ratios was used as the exon/intron ratio for the *APP* gene in a given cell. We also measured this ratio for two housekeeping genes (*GAPDH*, *ACTB*) and two source genes of germline pseudogene insertions (*SKA3* in AD3 and AD4, *ZNF100* in AD2) as negative and positive controls. One single cell (5087_MDA_02) showed a read-depth gain in the *APP* gene but it also showed the gain for many other brain-expressed genes including housekeeping genes, indicating genome-wide mRNA contamination. We thus excluded this single cell from further analysis.

To analyze exon-junction spanning reads, we utilized the same custom pipeline reported in the Lee study (https://github.com/christine-liu/exonjunction). We first remapped all our scWGS data to the human reference genome (GRCh38) using STAR (version 2.5.4a)[3] with the options described in the Lee study (--outSAMattributes All --outFilterScoreMinOverLread 0.8 --outSJfilterCountTotalMin 1 1 1 1). We then filtered the aligned BAM files using SAMtools (version 1.9)[15] to only contain the junction supporting reads by excluding the read with the tag 'jI:B:i,-1', as we described above. The filtered BAM files were converted into the BED12 format using bedtools bamtobed (version 2.27.1)[16]. Converted BED12 files were then applied to the custom pipeline to visualize reads spanning exon junctions of the *APP* gene. We obtained a few *APP* exon-junction-spanning reads from some scWGS data, however, we confirmed that all of them were the result of misalignment (*i.e.,* mis-split of the reads) by STAR due to short sequence homology between the intronic sequence of the exon-intron junction and the exonic sequence of the next intron-exon junction.

### Supplementary Figures

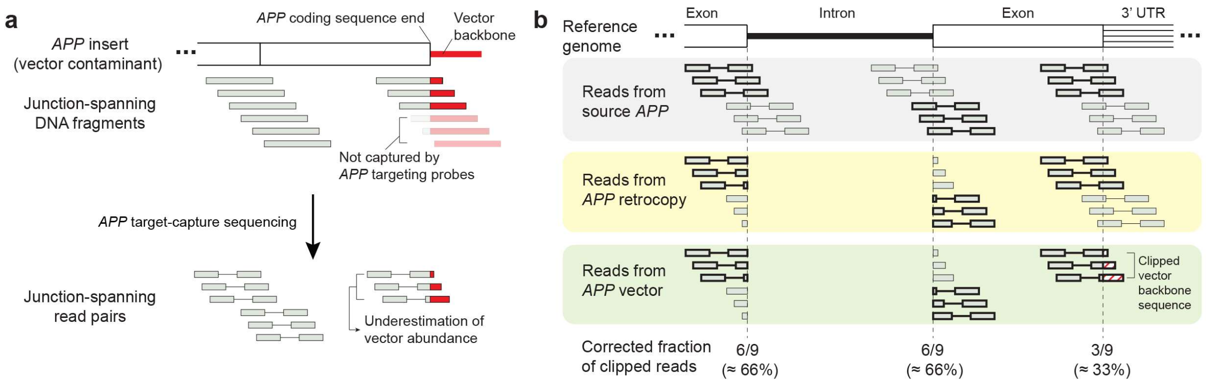


**Supplementary Fig. 1. Estimation of the fraction of vector inserts and true *APP* retrocopies from the targeted-capture sequencing data from the Lee study. a.** Difference in capturing efficiency between coding sequence ends and other exon junctions. In the targeted-capture data, DNA fragments largely consisting of vector backbone sequences would not be captured efficiently at coding sequence ends (shaded fragments). This difference results in the significant underestimation of the amount of vector contamination in the data. **b.** Adjustment of the bias in estimating clipped read fraction. To adjust for the difference in capturing efficiency, we considered the direction of the mate paired-end read, and used only the reads with mates mapped toward the inside of the exon to calculate the fraction (bolded reads). An average of the clipped read fractions of the two adjacent exon-intron junctions was used to represent the fraction of a given exon-exon junction.
